# The Genome of *Chenopodium ficifolium*: Developing Genetic Resources and a Diploid Model System for Allotetraploid Quinoa

**DOI:** 10.1101/2025.01.17.633571

**Authors:** Clayton D. Ludwig, Peter J. Maughan, Eric N. Jellen, Thomas M. Davis

## Abstract

High-quality nuclear, chloroplast, and preliminary mitochondrial genomes have been assembled and annotated for the B-genome diploid (BB: 2n = 2x = 18) figleaf goosefoot (*Chenopodium ficifolium*). The primary objective was to advance a simplified model system for genetic characterization and improvement of allotetraploid (AABB: 2n = 4x = 36) quinoa (*Chenopodium quinoa*), a nutritionally valuable, halophytic orphan crop. In addition to its diploidy and favorably small genome size, the *C. ficifolium* model provides a shorter generational period and smaller overall plant size as compared to *C. quinoa*, while displaying relevant agronomic trait variations amenable to gene-trait association studies. The *C. ficifolium* ‘Portsmouth’ nuclear genome was sequenced using PacBio HiFi Long Read technology and assembled using Hifiasm. After manual adjustments, the final ChenoFicP_1.0 assembly consisted of nine pseudochromosomes spanning 711.5 Mbp, while 22,617 genes were identified and annotated. BUSCO analyses indicated a nuclear genome completeness of 97.5%, and a proteome and transcriptome completeness of 98.4 percent. The chloroplast genome assembly detected two equally represented structural haplotypes differing in the orientation of the Short Single Copy region relative to the Long Single Copy region. Phylogenetic and parentage analyses pointed to an unspecified AA diploid species and away from *C. ficifolium* as the likely maternal chloroplast and mitochondrial genome donor(s) during the initial tetraploidization event in the *C. quinoa* lineage. Using the new ChenoFicP_1.0 reference genome, a GWAS was performed on a previously studied *C. ficifolium* F2 population to further define region(s) implicated in the control of three key agronomic traits: days to flowering, plant height, and branch number. This analysis localized control of all three traits to a 7 Mb interval on pseudochromosome Cf4. This region contains approximately 770 genes, including the *FTL1* locus, thus confirming and extending our prior, single-marker analysis showing association of these three traits with an *FTL1* amplicon length polymorphism. The use of these data to further develop *C. ficifolium* as a model species for genetics and breeding of quinoa serves to expand knowledge and germplasm resources for quinoa improvement.

## Introduction

*Chenopodium ficifolium* (figleaf goosefoot) is a weedy, Old-World B-genome diploid (BB: 2n = 2x = 18) relative of the cultivated allotetraploid (AABB: 2n = 4x = 36) pseudocereal crop quinoa (*Chenopodium quinoa* Willd.) (Walsh *et al*., 2015; Kolano *et al*., 2016) and its wild sister AABB tetraploid *Chenopodium berlandieri* (Walsh *et al*., 2015). The genus *Chenopodium* now resides in the family Amaranthaceae, its former Chenopodiaceae family home having recently been subsumed into the Amaranthaceae (Chase *et al*., 2016), which also hosts the relevant comparator *Beta vulgaris* (sugar beet). Putatively originating from Eurasia (Walsh *et al*., 2015), figleaf goosefoot is now widespread in North America, both as a weed of human disturbance and as a naturalized part of disturbed ecosystems. It is one of two recognized potential contributors, or a close relative thereof, of the B subgenome to the *C. quinoa* lineage during its initial tetraploidization event, the other B subgenome candidate being diploid *C. suecicum* (Jellen *et al*., 2011; Mandák *et al*., 2012; Walsh *et al*., 2015; Jarvis *et al*., 2017). Because figleaf goosefoot potentially contributed the B subgenome to quinoa, or is an extant sister species of the contributor, it is a highly relevant candidate to explore as a diploid model species in relation to quinoa’s genetic characterization and breeding, and its study may shed light on specific genes governing domestication-related traits in quinoa (Jellen *et al*., 2013; Walsh *et al*., 2015; Kolano *et al*., 2016; Subedi, Neff and Davis, 2021).

Absent a history of intensive breeding, *C. quinoa* has been categorized as an ‘orphan crop’ (Lemmon *et al*., 2018; Kumar and Bhalothia, 2020). However, quinoa is receiving increasing interest from consumers due to its high nutritional value and from agronomists and breeders due to its drought and salt tolerance (Gorinstein *et al*., 2002; Rojas *et al*., 2009; Vega-Gálvez *et al*., 2010; Jacobsen, 2011). These and other attributes make quinoa an attractive candidate for production in regions that cannot sustain the major grain crops. Furthermore, quinoa has the potential to serve as a valuable secondary crop, thereby contributing to food security. Thus, it is an attractive candidate for increased production, justifying investment in development of novel genetics and methods to streamline the breeding process (Jacobsen, 2003; Bazile, Jacobsen and Verniau, 2016).

Using *C. ficifolium* as a diploid model species has the potential to accelerate trait dissection and gene identification due to its comparative genetic simplicity; shorter generation time of ∼40 days compared to quinoa’s 90-120 days (Jacobsen, 1997); profuse flower, pollen, and seed production; and smaller plant size facilitating growth chamber cultivation (Subedi et al., 2021). Building upon prior, foundational studies (Štorchová *et al*., 2015, 2019), the targeted development of *C. ficifolium* as a diploid model species was initiated (Subedi, 2020; Subedi, Neff and Davis, 2021) with a focus on: i) producing intraspecific hybrids and segregating progenies; ii) investigating segregation in domestication-related traits of relevance to *C. quinoa* breeding programs; and iii) detecting gene-trait associations for key genes and traits.

In an F_2_ segregating population from an initial cross involving *C. ficifolium* accessions from Portsmouth (’P’) NH USA, and Quebec City (’QC’) Quebec Canada, allelic variation in the *Flowering Locus T-Like 1* (*FTL1*) marker locus was found to be associated with variation in days to flower, plant height, and branch number (Subedi, 2020; Subedi, Neff and Davis, 2021). Using previously designed primers (Cháb *et al*., 2008; Štorchová *et al*., 2015) to target and conveniently genotype an indel polymorphism within the *FTL1* gene, it was found that F2 plants homozygous for the ‘P’ allele flowered on average 12 days earlier than plants homozygous for the ‘QC’ allele, and two days earlier than heterozygotes (Subedi, 2020; Subedi, Neff and Davis, 2021). As a likely consequence of their decreased maturation time, ‘P’ allele homozygotes were also significantly shorter and had fewer branches than plants of the alternate genotypes. These results strongly implicated the *FTL1* chromosomal region in the control of three agronomically important traits, but did not rule out the possibility of influence from other genomic regions. While genotype-phenotype relationships involving *FTL1* must be further studied at both the diploid and the tetraploid level, this research demonstrates the potential value of *C. ficifolium* as a model system in which to accelerate discovery and characterization of gene-trait relationships. The goal of the work reported here is to expand knowledge and establish foundational genomic resources for the *C. ficifolium* model system.

## Materials and Methods

### Germplasm

All plant cultivation and phenotyping activities were conducted at the New Hampshire Agricultural Experiment Station (NHAES) at the University of New Hampshire (UNH). Two figleaf goosefoot accessions were employed in this study, both of which have been deposited as seed samples with the USDA North Central Plant Introduction Station (NCRPIS) at Ames Iowa. The ‘Portsmouth’ (’P’) accession (PI 698433) was collected by Erin Neff and Thomas Davis in Portsmouth, New Hampshire, USA (Neff, 2017), while the ‘Quebec City’ (’QC’) accession (PI 698434) was collected by Thomas Davis from Quebec City, Quebec, Canada. Crosses between the ‘P’ and ‘QC’ accessions to generate F_1_ and F_2_ populations, and confirmation of parentage using informative *FTL1* amplicons, were as previously described (Subedi, Neff and Davis, 2021). For purposes of comparison, we also employed in-house-numbered accessions 302-A of AA diploid *C. foggii*, and RB6 of AABB allotetraploid *C. berlandieri* var. *macrocalycium*, which had been collected by Erin Neff and Thomas Davis in southern Maine (Neff, 2017; Neff, Sullivan and Davis, 2018) and in Rye Beach New Hampshire, respectively; and *C. quinoa* accession QQ065 (PI 614880), obtained from USDA NPGS.

### DNA Isolation and Genome Sequencing

The figleaf goosefoot ‘Portsmouth’ (’P’) accession was used to construct the primary *de novo* reference genome assembly. Using leaf tissue provided by the UNH investigators, DNA from the ‘P’ accession was isolated at Brigham Young University (BYU) using methods designed to yield high molecular weight (HMW) DNA from plant tissues (Vaillancourt and Buell, 2019). The ‘P’ accession HMW DNA was sequenced at the BYU DNA Sequencing Center (DNASC) using the PacBio HiFi platform (Eid *et al*., 2009). In addition, this DNA was used at UNH to generate Nanopore sequence on an Oxford Nanopore Technologies (ONT) R9.4.1 flow cell using a library prepared with a rapid sequencing kit (SQK-RAD004) following methods described by Nanopore entitled “Rapid sequencing gDNA – whole genome amplification”.

Illumina sequence was generated at the UNH Hubbard Center for Genome Studies (HCGS) for the following: for the ‘P’ and ‘QC’ accessions of *C. ficifolium*; for 35 F2 generation individuals descended from a ‘P’ x ‘QC’ cross that was previously studied and described (Subedi et al., 2021); the RB6 accession of *C. berlandieri* var. *macrocalycium*; and for the 302-A accession of *C. foggii*. DNA for Illumina sequencing was isolated at UNH (Subedi et al., 2021) using a CTAB method described by Torres *et al*. (1993).

### Nuclear Genome Assembly

PacBio reads were assembled at BYU using Hifiasm Version v0.18.5-r499 (Cheng *et al*., 2021, 2022) with default settings. Subsequent correction, annotation, and analyses were conducted at UNH. Inspector (Chen *et al*., 2021) was used to detect and correct errors in the Hifiasm assembly followed by manual examination and adjustments, yielding the *C. ficifolium* Version 1.0 nuclear reference genome designated as ChenoFicP_1.0. Finally, BUSCO (Manni *et al*., 2021) and dependencies (Stanke *et al*., 2008; Camacho *et al*., 2009; Eddy, 2011; Mirarab, Nguyen and Warnow, 2012; Levy Karin, Mirdita and Söding, 2020; Li, 2023) using default settings and the embryophyta_odb10 database was used to calculate nuclear genome completeness. Telomeres were identified within pseudochromosomes (PCs) with the quarTeT toolkit program TeloExplorer (Lin *et al*., 2023) and within unplaced contigs using tidk (Brown, la Rosa and Mark, 2023). Centromeric regions were identified using the tool CentroMiner which is supplied with the quarTeT toolkit (Lin *et al*., 2023). Nuclear genome assembly data was visualized using Circos (Krzywinski *et al*., 2009) in combination with the deepStats toolkit (Richard, 2019) and BEDOPS (Neph *et al*., 2012) for track generation.

### Nuclear Genome Annotation

Once ChenoFicP_1.0 was finalized, repetitive elements were identified using RepeatModeler (Flynn *et al*., 2020). Those repetitive elements were then masked using RepeatMasker (Tarailo-graovac and Chen, 2009; Flynn *et al*., 2020; Storer *et al*., 2021) as the first step in an annotation pipeline patterned after Card *et al*. (2019). The RepeatMasker scripts calcDivergenceFromAlign.pl and createRepeatLandscape.pl were used to analyze output files, determine repetitive element content, and produce data visualizations. A BRAKER IsoSeq compatible singularity container (Bruuna, Gabriel and Hoff, 2024) was passed the fasta file containing soft-masked repetitive elements and Pacific Biosciences IsoSeq data aligned to the genome using minimap2 (Li, 2018). BRAKER flags were set to use the Viridiplantae OrthoDB database (Kuznetsov *et al*., 2023) for protein sequence, and the Embryophyta odb10 database (Manni *et al*., 2021) as the Busco lineage.

The resulting GFF3 annotation file was passed to MAKER (Holt and Yandell, 2011) as a prediction GFF file, as were the cds-transcripts and proteins files which were used for EST evidence and protein homology evidence, respectively. MAKER was used to update annotations and to provide Annotation Edit Distance (AED) scores. AED values for the final annotation file were produced using AED_cdf_generator.pl (https://github.com/mscampbell). Once complete, the output files for each pseudochromosome were merged to create one contiguous file. This output file contained gene predictions, however without functional annotations. To integrate functional details and GO terms, BLASTP (Johnson *et al*., 2008) using the UniProtKB/Swiss-Prot database (Bateman *et al*., 2015), as well as InterProScan (Jones *et al*., 2014) were passed the protein file produced by MAKER, and the resulting information was integrated into a single file using the maker_functional_gff and ipr_update_gff commands, and sorted with AGAT (Dainat, Hereñú and Pucholt, 2020), which yielded the final GFF3 annotation file.

A genome-wide synteny analysis was performed comparing the ChenoFicP_1.0 assembly to the B subgenome of the *C. quinoa* V2 reference genome (Rey *et al*., 2023) and to the *Beta vulgaris* EL10.1 reference genome (McGrath *et al*., 2023). As a co-member of the Amaranthaceae family, diploid (2n = 2x = 18) *B. vulgaris* (sugar beet) has been used in previous genomic comparisons with quinoa (Jarvis *et al*., 2017; Rey *et al*., 2023). To produce the synteny analysis, *C. quinoa* annotations and protein fasta files were downloaded from CoGe (id60716), and *B. vulgaris* files from NCBI (GCF_002917755.1). Annotation files were converted to BED format via agat_convert_sp_gff2bed.pl (Dainat, Hereñú and Pucholt, 2020), before being manually rearranged to comply with requirements for MCScanX_h (Wang *et al*., 2012). BlastP was used to compare the proteome of *C. ficifolium* with those of *B. vulgaris* and *C. quinoa* (Camacho *et al*., 2009), yielding two blast output files which were merged and passed to MCScanX_h, along with BED files containing positional information for the genes in the blast files. For this analysis, MCScanX used default filtering parameters. SynVisio (Bandi and Gutwin, 2020) was used to visualize the resulting files, illuminating rearrangements and structural variations.

### Chloroplast Genome Assembly and Annotation

The ‘P’ chloroplast genome was initially assembled from extracted PacBio reads using GetOrganelle v1.7.6.1 (Jin *et al*., 2020) with dependencies SPAdes (Bankevich *et al*., 2012), Bowtie2 (Langmead and Salzberg, 2012), BLAST+ (Camacho *et al*., 2009), and Bandage (Wick *et al*., 2015), and annotated with the online tool GeSeq (Tillich *et al*., 2017). GetOrganelle was called using get_organelle_from_reads.py and was supplied with raw PacBio reads using the unpaired ‘-u’ flag. Embplant_pt was selected as the genome type, and the target genome size was set to 150 kb which is the approximate size of the *C. quinoa* chloroplast genome (Hong *et al*., 2017). For the purpose of determining the relative abundance of each detected haplotype of the ‘P’ chloroplast genome based on the numbers of supporting Nanopore reads, Cp-hap (Wang, Lanfear and Gaut, 2019) was passed reads produced via an ONT R9.4.1 flow cell as previously noted. The final output file produced from GetOrganelle was passed to GeSeq (https://chlorobox.mpimp-golm.mpg.de/geseq.html) (Tillich *et al*., 2017) for visualization and annotation.

### Mitochondrial Genome Assembly and Annotation

A contig from the Hifiasm assembly was identified as mitochondrial by aligning all unplaced contigs to the quinoa mitochondrial reference sequence NC_041093.1 (Maughan *et al*., 2019) with BWA MEM (Li and Durbin, 2009), and manually selecting a long, well-aligned contig. This contig was used as a seed sequence and passed to a *de novo* organelle assembly pipeline described by Jarvis et al., (2022) to assemble the mitochondrial genome. The pipeline was used in two iterations, initially using the HiFi dataset which produced a contig that was passed to the mitochondrial assembly pipeline in conjunction with the long read Nanopore dataset. The Nanopore dataset was then aligned to this contig using minimap2 (Li, 2018), reads matching GC content and length thresholds were stripped from the alignment file via SeqKit (Shen *et al*., 2016), and these reads were passed to Canu (Koren *et al*., 2017) for *de novo* assembly. Nanopore reads were later used to resolve assembly structure via comparing alignments to the draft mitogenome assembly. The output contig from Canu was manually circularized by locating regions at the start and end of the file that shared 100 percent sequence identity. The circularized version of the assembly was polished using NextPolish (Hu *et al*., 2020) and the PacBio read set to correct single nucleotide errors. Annotation was performed by GeSeq. Importantly, GeSeq was set to pass third party mitochondrial NCBI RefSeqs from both *C. quinoa* (NC_041093.1) and *Beta vulgaris* subsp. *vulgaris* (NC_002511.2) to BLAT (Kent, 2002) which used annotations from those genomes to locate and annotate homologs within the *C. ficifolium* mitogenome.

### Chloroplast and Mitochondrial Inheritance Patterns in *C. ficifolium*

Chloroplast and mitochondrial reference-guided assemblies for ‘QC’, one F1 individual, and four F2 individuals, were constructed and aligned to the respective ‘P’ organelle genomes using BWA mem (Li and Durbin, 2009) to identify heritable organelle polymorphisms differentiating the two parents. The resulting SAM files were then converted to BAM, sorted, indexed, and a consensus was called using the respective samtools modules (Danecek *et al*., 2021). Once BAM files for each individual were converted to fasta, freebayes-parallel (Garrison and Marth, 2012; Tange, 2018) was used to call variants. Freebayes-parallel was invoked with options to produce a gVCF file, include genotype qualities, use best n alleles set to 4, and cap read depth to 5000. VCF files were filtered to remove low quality calls using vcffilter from vcflib (Garrison *et al*., 2022) and bcftools (Danecek *et al*., 2021). Sites or calls were pruned from the chloroplast multisample vcf file if: a) the average depth across all samples was below 100; b) or the site quality was below 998; c) or the genotype quality phred score was below 30; d) or depth for a given sample was below 50; e) or more than half of individuals were not represented; f) or the allele balance was below 25 percent or greater than 75 percent; g) or there was a discrepancy between the number of paired and unpaired reads aligned. The vcf file generated for the mitochondrial genome using the same samples was filtered using the same criteria with the addition of a cap of 3000 on the average depth per sample. The filtering criteria and script were based on those provided by http://www.ddocent.com/filtering/. The resulting filtered vcf files were visualized with IGV (Thorvaldsdóttir, Robinson and Mesirov, 2013).

### Chloroplast and Mitochondrial Genome Ancestries

To illuminate the possible role of *C. ficifolium* in the cpDNA ancestries of allotetraploids *C. quinoa* and *C. berlandieri*, a minimal phylogenetic tree was generated using six chloroplast genome (cpDNA) assemblies representing four *Chenopodium* species and including beet (*Beta vulgaris*) as an outgroup. The *C. quinoa* and *B. vulgaris* chloroplast genomes used were NCBI accessions MK159176.1 and OU343016.1, respectively. All other accessions included in this analysis were assembled by us *de novo* from raw reads using GetOrganelle (Jin *et al*., 2020). We assembled the *C. berlandieri* chloroplast genome from Illumina 2×150 PE reads previously uploaded to NCBI by Mark Samuels from the University of Montreal (Accession: PRJNA895488) (Samuels *et al*., 2023). The *C. ficifolium* ‘P’ chloroplast genome was *de novo* assembled using sequence data extracted from the PacBio dataset as described above, while the *C. ficifolium* ‘QC’ and the *C. foggii* 302-A chloroplast genomes were assembled from Illumina data produced at the UNH-HCGS. These assemblies were aligned using Mafft (Katoh and Standley, 2013) with default (auto) settings, and a tree was produced using RAxML V. 8.2.12 with the GTRCAT model (Stamatakis, 2014). Trees were visualized with TreeViewer (Bianchini and Sánchez-Baracaldo, 2023).

For a parallel purpose, a minimal mitochondrial phylogeny was produced by aligning raw reads, either generated by the HCGS for *C. foggii* 302-A, and ‘P’ and ‘QC’ *C. ficifolium* accessions or downloaded from NCBI, to the quinoa mitochondrial reference genome NC_041093.1 (Maughan *et al*., 2019). *B. vulgaris* (SRR6305245) (McGrath *et al*., 2023) and *C. berlandieri* (PRJNA895488) (Samuels *et al*., 2023) read sets were downloaded from NCBI. All read data used in this analysis were trimmed and cleaned with Trimmomatic (Bolger, Lohse and Usadel, 2014) using the primer sequence file “TruSeq3-PE-2.fa” invoked with the following settings: ILLUMINACLIP:TruSeq3-PE-2.fa:2:30:10 LEADING:3 TRAILING:3 SLIDINGWINDOW:4:15 MINLEN:36. FastQC (https://www.bioinformatics.babraham.ac.uk/projects/fastqc/) was used to visualize and validate the Trimmomatic output. These reads were then aligned to the quinoa mitochondrial reference genome (Maughan *et al*., 2019) using BWA mem (Li and Durbin, 2009). Using Samtools view, sort, index, and consensus (Danecek *et al*., 2021), the BWA produced SAM file was converted to a fasta file containing the consensus sequence. These six fasta files, those produced from reads plus the quinoa mitogenome file used as a reference in the alignment process, were merged into a multi-fasta and Mafft (Katoh and Standley, 2013) was used with default (auto) settings to produce an MSA. A tree was produced using RAxML V. 8.2.12 (Stamatakis, 2014) with the GTRCAT model and *B. vulgaris* specified as the outgroup.

### Genome Wide Association Studies

A subset of 21 ‘P’ x ‘QC’ F2 individuals chosen from the 35 sequenced at the HCGS on the basis of high read depth were used in a GWAS analysis to locate regions throughout the genome implicated in the control of branch angle, internode length, flowering time, plant height, and number of lateral branches. BWA MEM was used to align reads from F2 individuals to the *C. ficifolium* ‘P’ V1.0 reference genome. Resulting sam files were converted to bam, sorted, and indexed via samtools (Danecek *et al*., 2021), and read depths visualized with WGSCoveragePlotter (Lindenbaum, 2015). GATK AddOrReplaceReadGroups (Van der Auwera and O’Connor, 2020) was called to change read group IDs to reflect the sample name to make variant calling possible. Freebayes-parallel (Garrison and Marth, 2012; Tange, 2018) was used to generate a vcf file containing both InDel and SNP variants for all 35 initial individuals. The resulting vcf file was filtered to remove low quality calls following guidelines from http://www.ddocent.com/filtering/. The filtered vcf data in conjunction with phenotypic data collected by Subedi (2020) (Supplemental Table 1), and name and positional data extracted from the GFF3 annotation file were supplied to the package vcf2gwas (Vogt, Shirsekar and Weigel, 2022) which produced Manhattan plots and positional data for putatively causative SNPs.

A region of interest on Cf4 identified via vcf2gwas was then closely examined with respect to gene content and local synteny and collinearity with the *C. quinoa* V2 assembly (Rey *et al*., 2023). For this purpose, this approximately 7 Mb region of interest was manually excised from the full annotation file associated with our ChenoFicP_1.0 assembly, Revigo (Supek *et al*., 2011) was used to analyze pathways and relationships of genes with attached GO terms, and gene IDs were used to subset the proteome fasta file. BlastP and MCScanX were used to compare this protein subset to the complete proteome of *C. quinoa* and create a homology file. This file, plus the required concatenated BED files were passed to MCScanX_h, which used default filtering criteria with the exception of MATCH_SIZE, which was set to 9 to only include matches with 10 or more collinear genes. The MCScanX pipeline developed by BDX consulting (Wang *et al*., 2024) was used to convert data to a format compatible with Circos (Krzywinski *et al*., 2009).

## Results

### Nuclear Genome Assembly

The PacBio HiFi read set from *C. ficifolium* ‘P’ comprised a total of 26.5 Gb of sequence with an N50 of ∼14 Kb. These data were supplied to Hifiasm Version 18, yielding 781 contigs of cumulative length 759,656,873 bp (∼760 Mb) with an average read depth of 34.8x (Table 1).

**Table 1.**
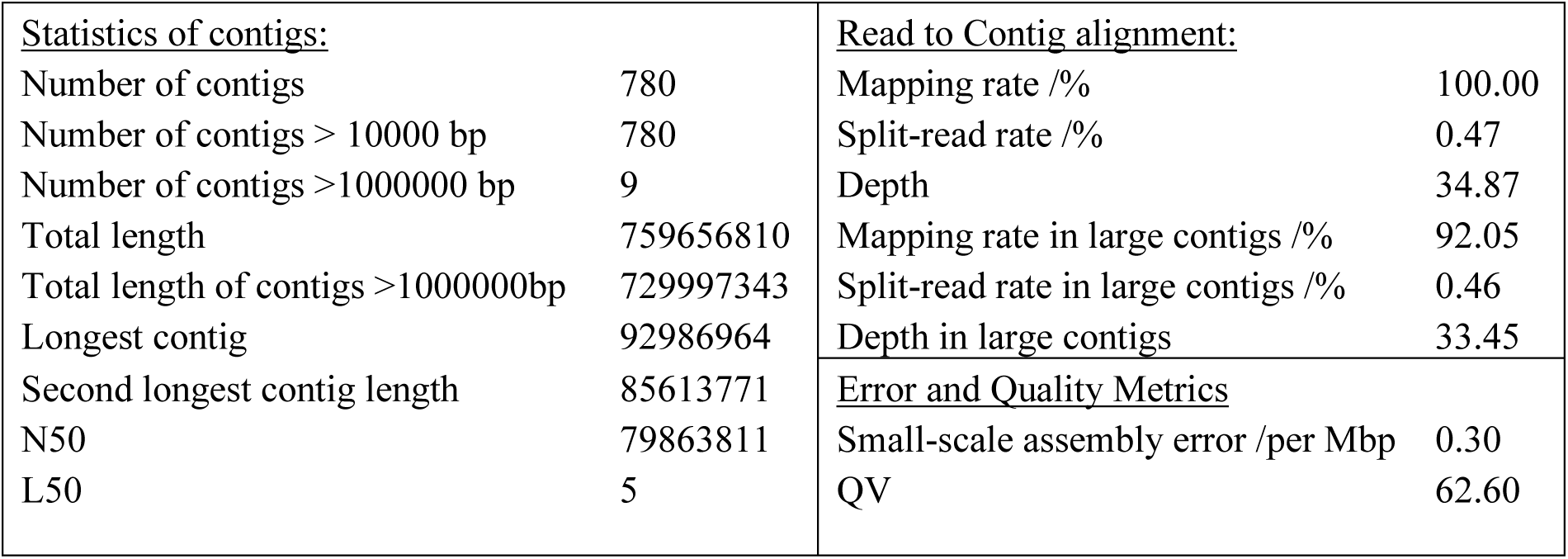
Hifiasm assembly statistics.

| <u>Statistics of contigs:</u> |  | <u>Read to Contig alignment:</u> |  |
| --- | --- | --- | --- |
| Number of contigs | 780 | Mapping rate /% | 100.00 |
| Number of contigs > 10000 bp | 780 | Split-read rate /% | 0.47 |
| Number of contigs >1000000 bp | 9 | Depth | 34.87 |
| Total length | 759656810 | Mapping rate in large contigs /% | 92.05 |
| Total length of contigs >1000000bp | 729997343 | Split-read rate in large contigs /% | 0.46 |
| Longest contig | 92986964 | Depth in large contigs | 33.45 |
| Second longest contig length | 85613771 | <u>Error and Quality Metrics</u> |  |
| N50 | 79863811 | Small-scale assembly error /per Mbp | 0.30 |
| L50 | 5 | QV | 62.60 |

Of these contigs, the largest nine ranged in size from a maximum of 92,986,964 bp down to 70,697,261 bp, while the tenth largest (contig ptg000009l) was 8,313,269 bp (Table 2). The remaining 771 contigs were all much smaller: less than 1 Mbp. On the basis of these size discontinuities, and the prior expectation based on the *C. ficifolium* chromosome number (n = x = 9) (Walsh *et al*., 2015), the nine largest Hifiasm contigs were defined as comprising the initial pseudochromosome (PC) assembly. These nine PCs were then numbered and oriented to maximize correspondence with the *Beta vulgaris* EL10.1 reference genome (McGrath *et al*., 2023), as was done previously in the assembly of the quinoa Version 1 assembly (Jarvis *et al*., 2017). Inspector subsequently identified and corrected three structural errors, and 228 small-scale assembly errors, reporting a QV score of 63.39 after correction for this assembly (Figure 1). The G/C content of this genome is 36.9 percent, and varies between approximately 20 and 80 percent per 100 Kb window (Figure 1 Ring D). G/C rich regions are often centrally located within pseudochromosomes; however, centromeric regions on the whole have an average G/C content, with pseudochromosome Cf1 having the highest content with 45.47 percent, and the average across all centromeres being 37.75 percent. When comparing the *C. ficifolium* ‘QC’ accession to the *C. ficifolium* ‘P’ assembly (Figure 1 Ring C), the number of SNPs differentiating the two per 100 Kb window ranges from as low as 12 to a maximum of 2,321, or approximately one SNP per 43 bases, as found on PC Cf1 from base 11,800,000 to base 11,900,000.

**Figure 1.**
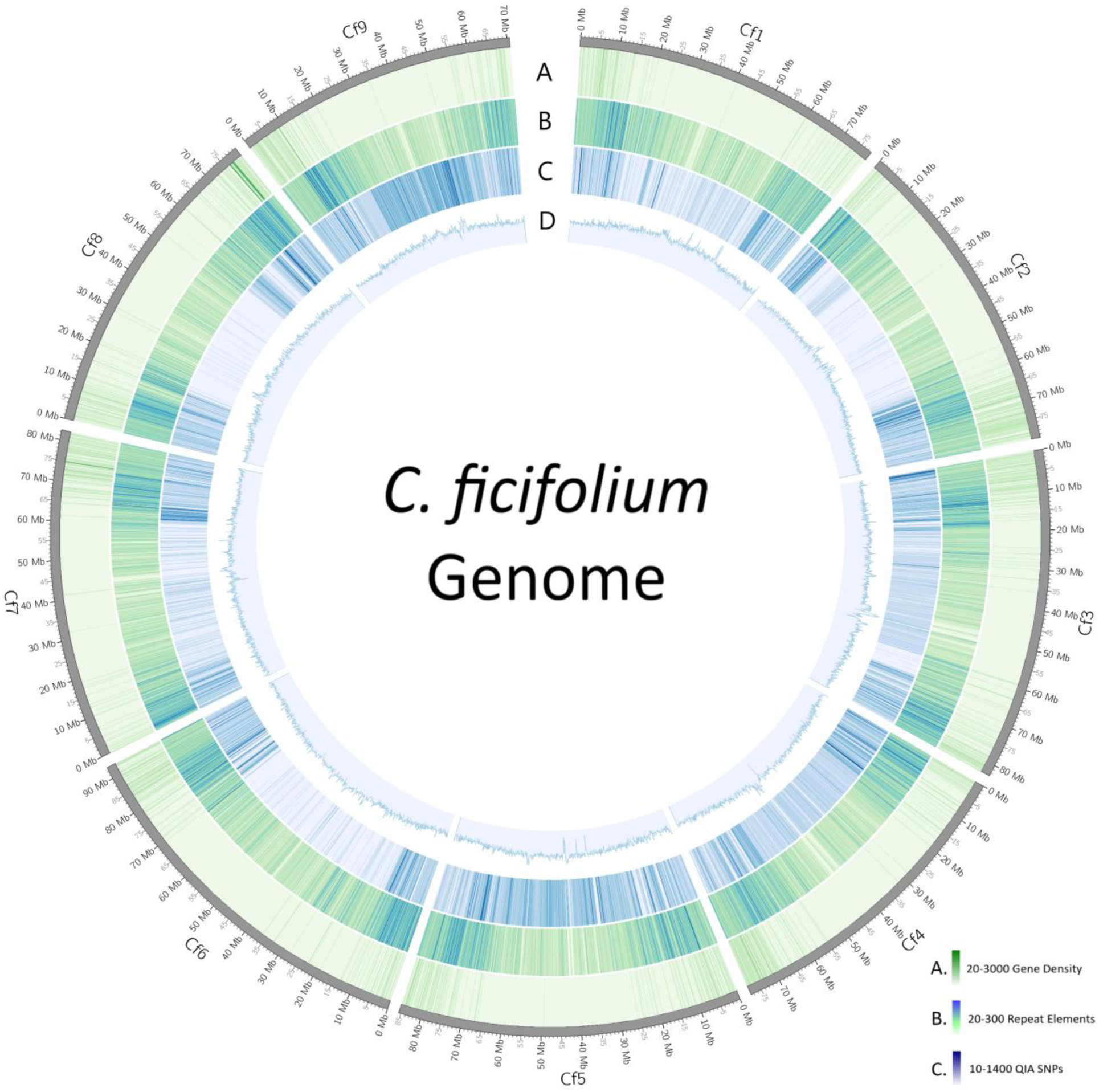
This assembly of the C. ficifolium ‘UNH_ChenoFicP_1.0’ nuclear genome comprises nine pseudochromosomes and spans 729.9 Mb. **A.** Gene density ranging from 20 to 3000 genes represented per 100 kb window. **B.** Repetitive element density ranging from 20 to 300 elements per 100 kb window **C.** SNPs present compared to C. ficifolium accession ‘QC’ per 100 kb window. **D.** G/C content. Graph represents 20 - 55 percent G/C.

**Table 2.**
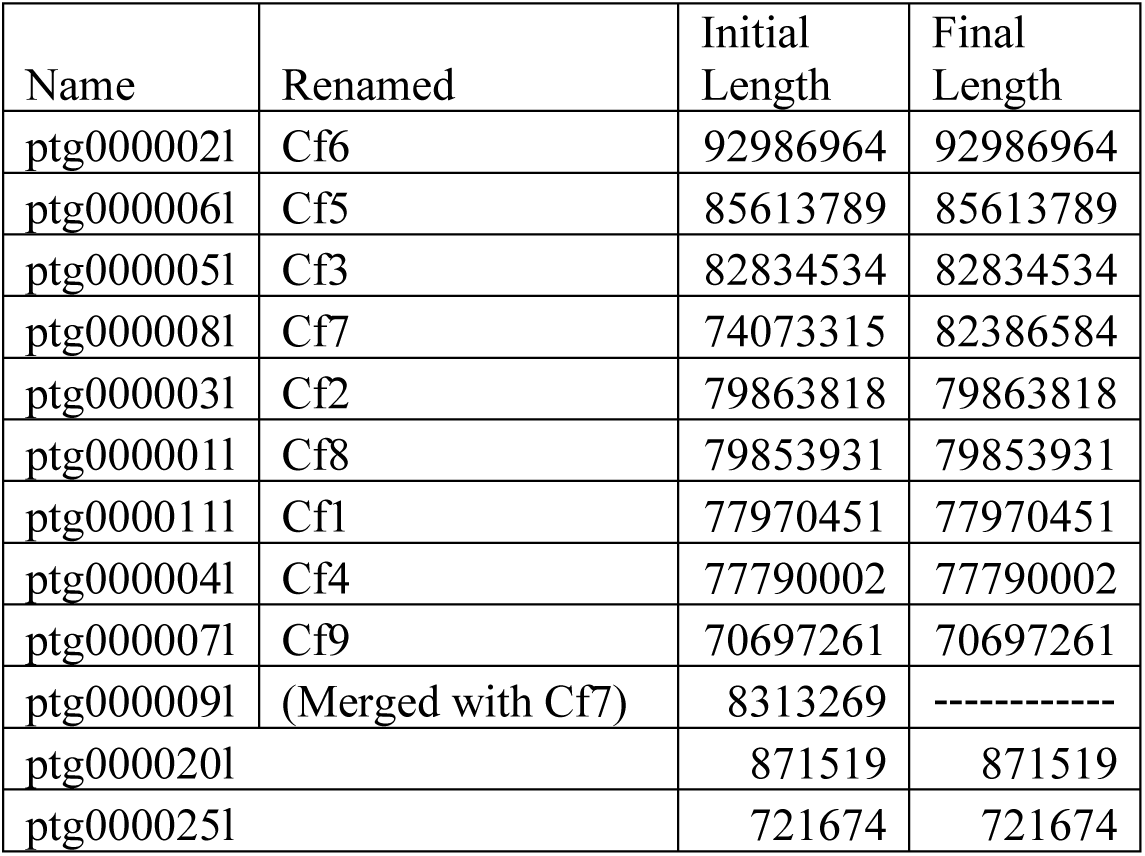
Original contig names, pseudochromosome designations, and contig lengths of the primary assembly plus the two largest unplaced contigs after ptg000009l were appended to Cf7.

The initial Hifiasm assembly was telomere-to-telomere for PCs Cf3, Cf4, Cf5, Cf6, Cf8, and Cf9, with Cf1 and Cf2 having only one telomere each, and Cf7 having a partially assembled telomere which was oriented in a backwards direction. A homology search of contig ptg000009l, the only unplaced contig larger than 1 Mbp, relative to the quinoa B subgenome (Rey *et al*., 2023) revealed that this contig was syntenic to the distal end of quinoa chromosome Cq7B, including a telomeric sequence, and thus directed its placement onto Cf7. Following this adjustment, the final “ChenoFicP_1.0” assembly of nine pseudochromosomes had a total length of 729,997,451 bp (∼730 Mb). The CentroMiner package identified one region per chromosome as being centromeric (Supplemental Table 2) and produced positional data. Centromeric and telomeric positional data are provided in Supplemental Table 2. BUSCO completeness of the reference genome without unplaced contigs is 96.7 percent, and with unplaced contigs is 97.2 percent (Figure 2B).

**Figure 2.**
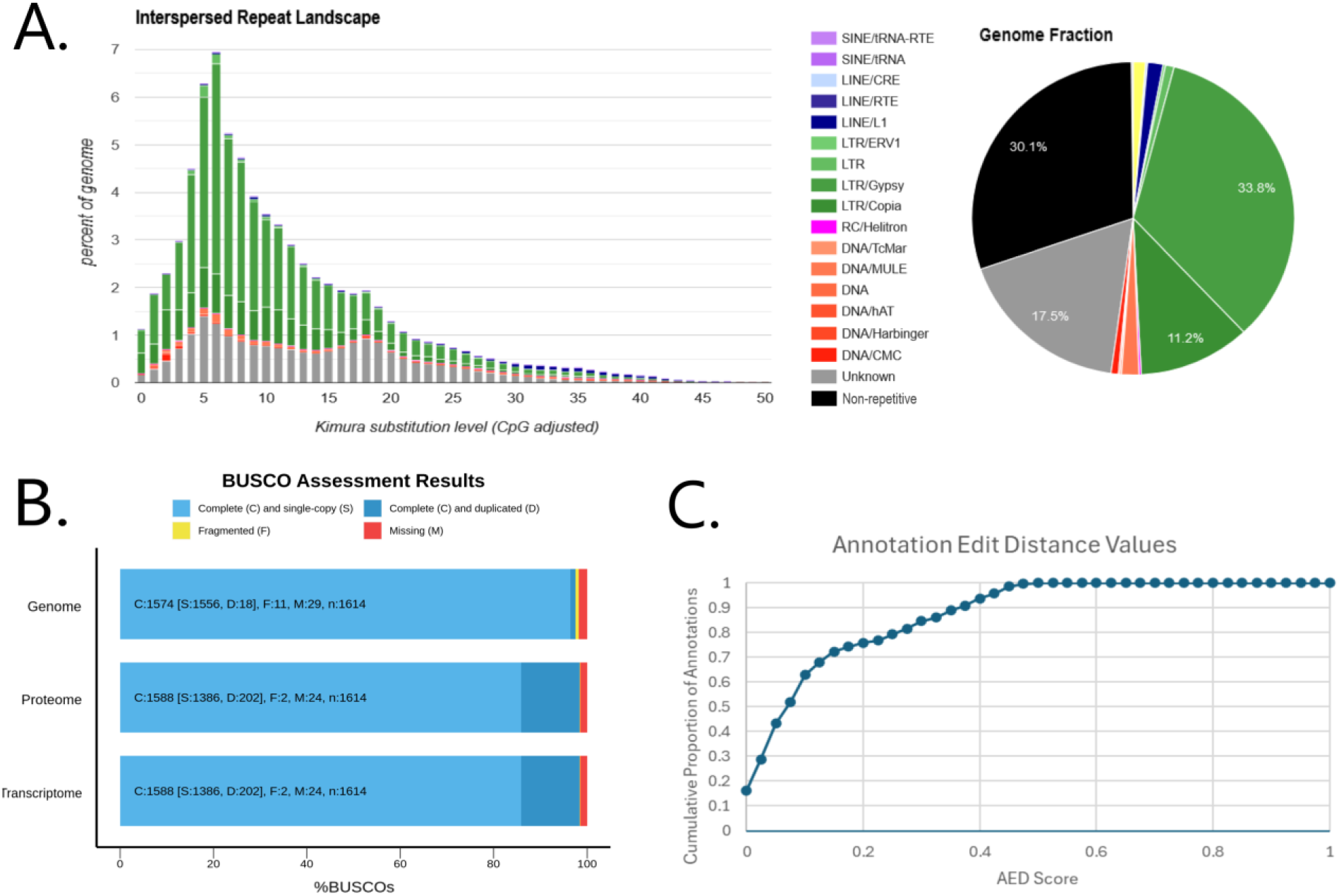
**A**. Repetitive element analysis and summary graph produced by RepeatModeler. Green: LTR elements, Grey: Unknown repetitive elements, Black: Non-repetitive sequence. **B**. BUSCO analyses of completeness for the masked genome used in the annotation process (top), the proteome (middle), and transcriptome (bottom). **C**. AED (annotation edit distance) scores of gene reannotations produced by Maker. Lower AED values correspond with higher levels of agreement between the annotation and evidence, with 0 representing perfect agreement between the two.

The remaining 771 unplaced contigs comprised a total of about 29.6 Mb, of which 594 contigs (19.6 Mb) and 6 contigs (626 Kb) were subsequently assigned on the basis of sequence homology to the chloroplast and mitochondrial genomes, respectively, leaving a final total of 171 contigs (9.4 Mb) unplaced. Further investigation into unplaced contigs using tidk indicated that contigs ptg000020 (871,519 bp) and ptg000025 (721,674 bp) both contain telomeric sequence. These contigs were found by BWA to likely belong on the upstream ends of Cf2 and Cf7 respectively; however, they have not been included in the ChenoFicP_1.0 assembly pending further confirmation based upon anticipated long read data.

### Repetitive Element Identification and Genome Annotation

Data produced by RepeatModeler indicates that 69.9 percent of the ChenoFicP_1.0 genome is comprised of repetitive elements, with the largest contributors being LTR/Gypsy (33.8 percent) and LTR/Copia (11.2 percent) type repetitive sequences, while unknown repeats account for 17.5 percent of the total genome fraction. These repetitive elements tend to cluster towards telomeres in (Figure 1 Ring B). Non-repetitive sequence accounts for 30.1 percent of the genome (Figure 2A).

The final annotation file was used to calculate relative gene density (Figure 1 Ring A). The BUSCO analysis yielded a transcriptome and proteome completeness of 98.4 percent (Figure 2B). Resultant AED values for the final annotation are graphed in Figure 2C. In total, 22,617 genes were identified and annotated with the average gene length (including introns and UTRs, as defined by Maker utilizing unprocessed mRNA transcripts) being 3,958 bp.

### Synteny with Quinoa B-subgenome Chromosomes

The MCScanX synteny analysis of the ChenoFicP_1.0 assembly relative to *B. vulgaris* EL10.1 (McGrath *et al*., 2023) and the quinoa V2 assembly B-subgenome (Rey *et al*., 2023) revealed numerous structural variations. In both comparisons, translocational differences tended to involve small distal regions near telomeres. Large inversional differences were revealed in ChenoFicP_1.0 chromosomes Cf1, Cf3, and Cf4 relative to the quinoa B-subgenome homologues, the largest of which was on Cf3 and spanned approximately 63.2 Mb (Figure 3). These inversions were positioned centrally, and in every case for chromosomes Cf1, Cf3, and Cf4 were pericentric.

**Figure 3.**
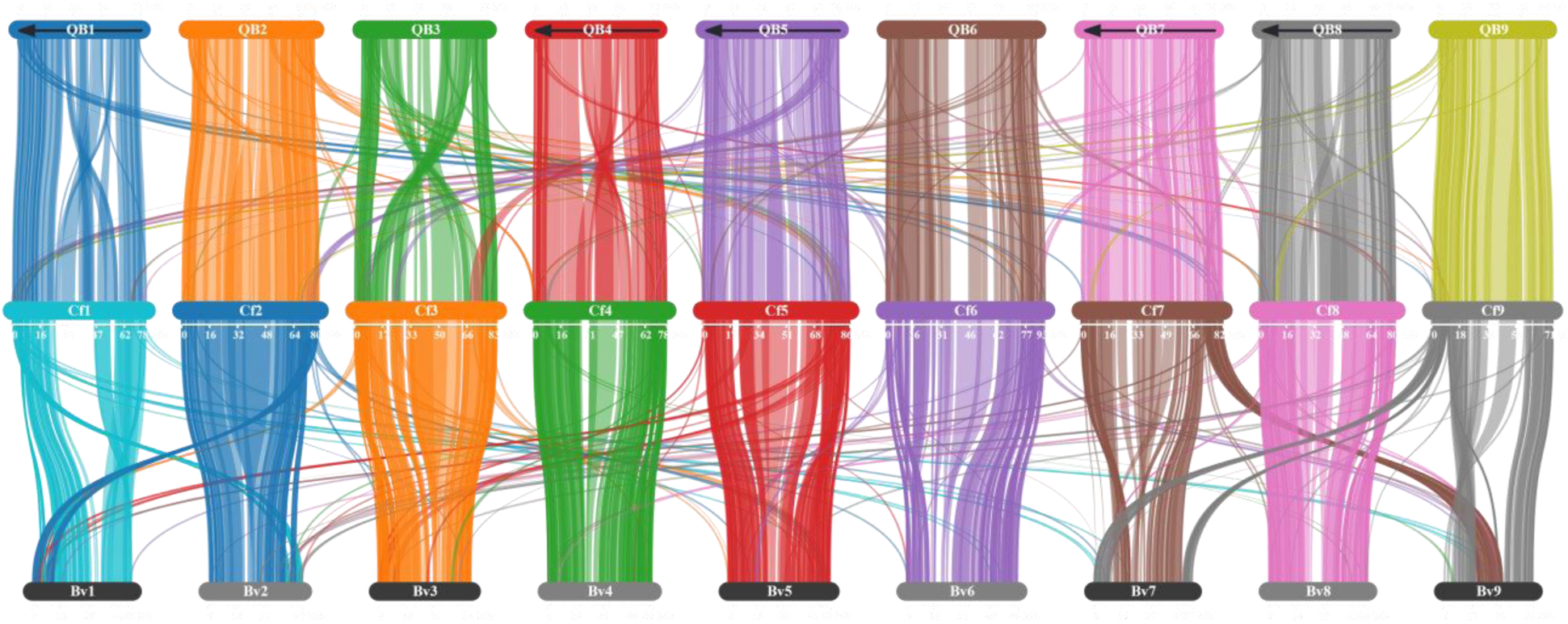
Synteny conservation diagram comparing the assembled pseudochromosomes of C. ficifolium (center) to the B subgenome of quinoa (top), and B. vulgaris EL10.1 (bottom). Chromosomes 1, 4, 5, 7, and 8 of the quinoa B subgenome were manually reoriented (flipped – indicated by arrows) to match those of C. ficifolium for illustrative purposes.

### Chloroplast Genome Assembly

GetOrganelle produced two versions of the *C. ficifolium* chloroplast genome assembly, termed “Path 1” and “Path 2”, both 151,826 bp in length, but differing in their orientations of the Short Single Copy (SSC) region relative to the Long Single Copy (LSC) region (Figure 4A). When supplied with approximately 263 Mb of Nanopore reads, Cp-hap (Wang, Lanfear and Gaut, 2019) returned approximately equal long read support for both Path 1 (248 reads) and Path 2 (242 reads) orientations, where the Path 2 orientation (Figure 4A, lower) coincided with that of the quinoa chloroplast assembly (MK159176.1) reported by Maughan et al. (2019).

**Figure 4.**
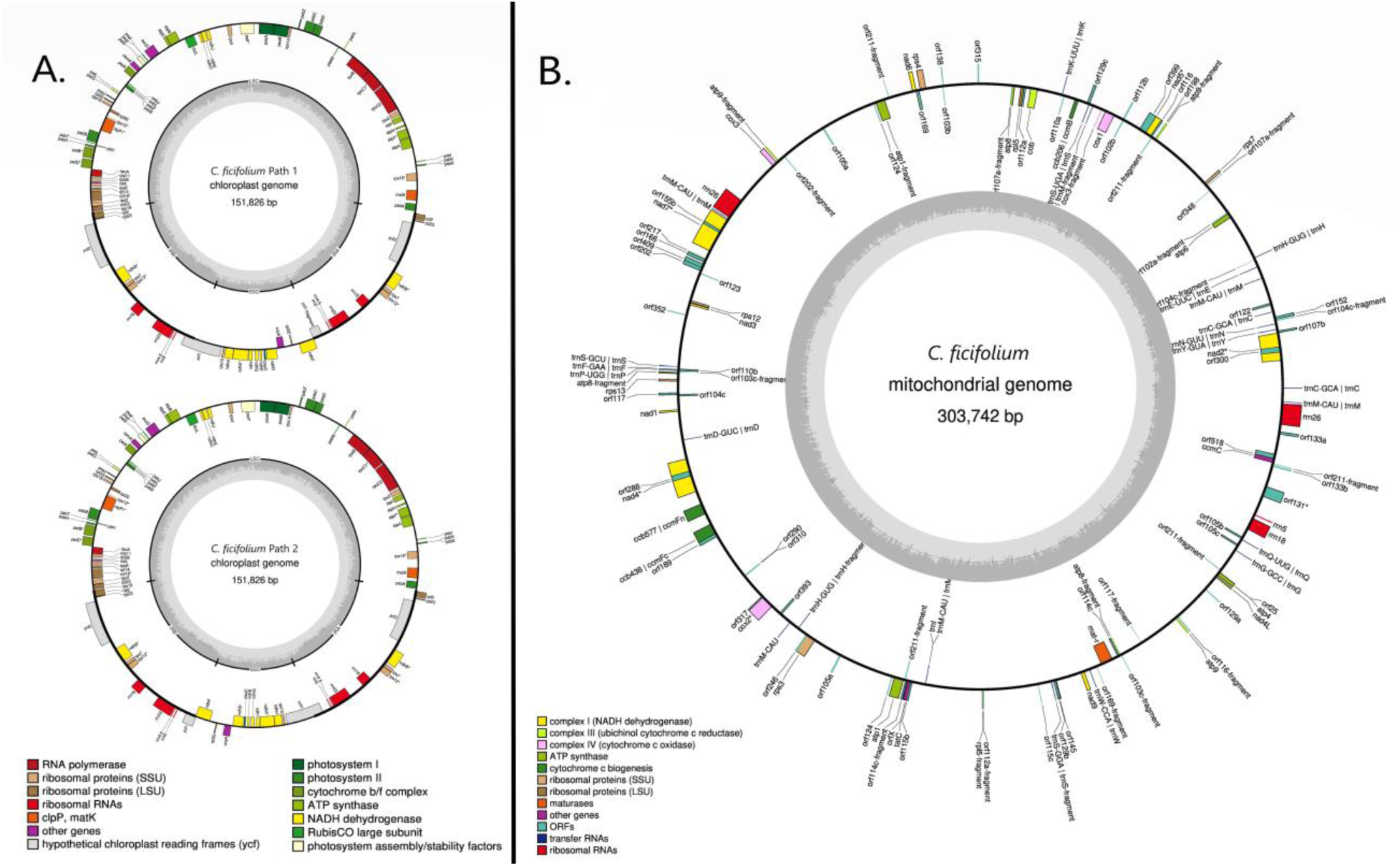
**A**. Both assembly versions of the C. ficifolium chloroplast genome represented by ‘Path 1’ and ‘Path 2’. These assemblies differ only in the direction of the Short Single Copy region. ‘Path 2’ (lower) was concordant with the C. quinoa chloroplast genome (MK159176). Genes inside the circle are transcribed in the clockwise direction, genes outside are transcribed counterclockwise. **B**. Gene map of the C. ficifolium mitogenome. The mitochondrial genome contains 118 protein coding genes, 23 tRNA genes, and is ∼304 Kb in length.

In both Path 1 and Path 2 orientations, the large single copy region is 83,900 bp, the short single copy is 18,168 bp, and each of the inverted repeat regions are 15,109 bases in length. The chloroplast genome regional boundaries are specified in Table 3. The assembly, including both inverted repeat regions, has a GC content of 37.3 percent and comprises 131 genes, of which there are 86 protein-encoding, 37 tRNA, and 8 rRNA genes. For comparison, the previously published (Kim et al., 2019) *C. ficifolium* chloroplast genome assembly (MK182725) had 129 genes, including 84 protein-encoding, 37 tRNA, and 8 rRNA.

**Table 3.** Start and end positions of the four regions of the chloroplast assembly. Note that these positions apply to both Path 1 and Path 2 haplotypes; however, the SSC region is inverted in Path 1 relative to Path 2.

| Cp Region | Start | End | No. Genes | Protein Encoding |
| --- | --- | --- | --- | --- |
| LSC | 1 | 83670 | 85 | 63 |
| IR | 83671 | 108779 | 16 | 5 |
| SSC | 108780 | 126717 | 14 | 13 |
| IR | 126718 | 151826 | 16 | 5 |
|  |  | Total | 131 | 86 |

### Mitochondrial Genome Assembly

A 275 kb contig containing a partially assembled mitochondrial genome from the ‘P’ accession was initially identified from a preliminary Hifiasm assembly. In an iterative sequence seeding process, Nanopore reads were aligned to this contig, extracted from the resulting alignment, assembled *de novo*, and reads were once again aligned. After two rounds of this process, a contig was produced that was manually circularized, and polished with NextPolish using HiFi reads. The resulting ‘P’ mitochondrial genome assembly has a GC content of 43.8 percent and comprises 90 genes, of which 33 are protein-coding (Figure 4B). The *C. ficifolium* mitochondrial genome assembly is similar in length to that of *C. quinoa* (MK182703), at 303,742 bp versus 319,505 bp respectively, and in protein-coding genes, with 33 and 30 respectively (Maughan *et al*., 2019), although these two assemblies display several structural differences.

### Organelle Genomes’ Modes of Transmission

When plastid genome mode of transmission was examined using multisample vcf files containing both parents, ‘P’ and ‘QC’, an F1, and four F2 individuals, two distinct sequence haplotypes were revealed. Of the respective polymorphisms, two were single nucleotide indels, while the remaining 17 were SNPs (Supplemental Table 3). The F1 and all F2 individuals share the same sequence haplotype as the maternal parent (’P’) as distinct from the ‘QC’ paternal haplotype (Figure 5A), thereby demonstrating maternal transmission of the chloroplast genome in the ‘P’ x ‘QC’ cross used to generate the respective F1 and F2 progenies.

**Figure 5.**
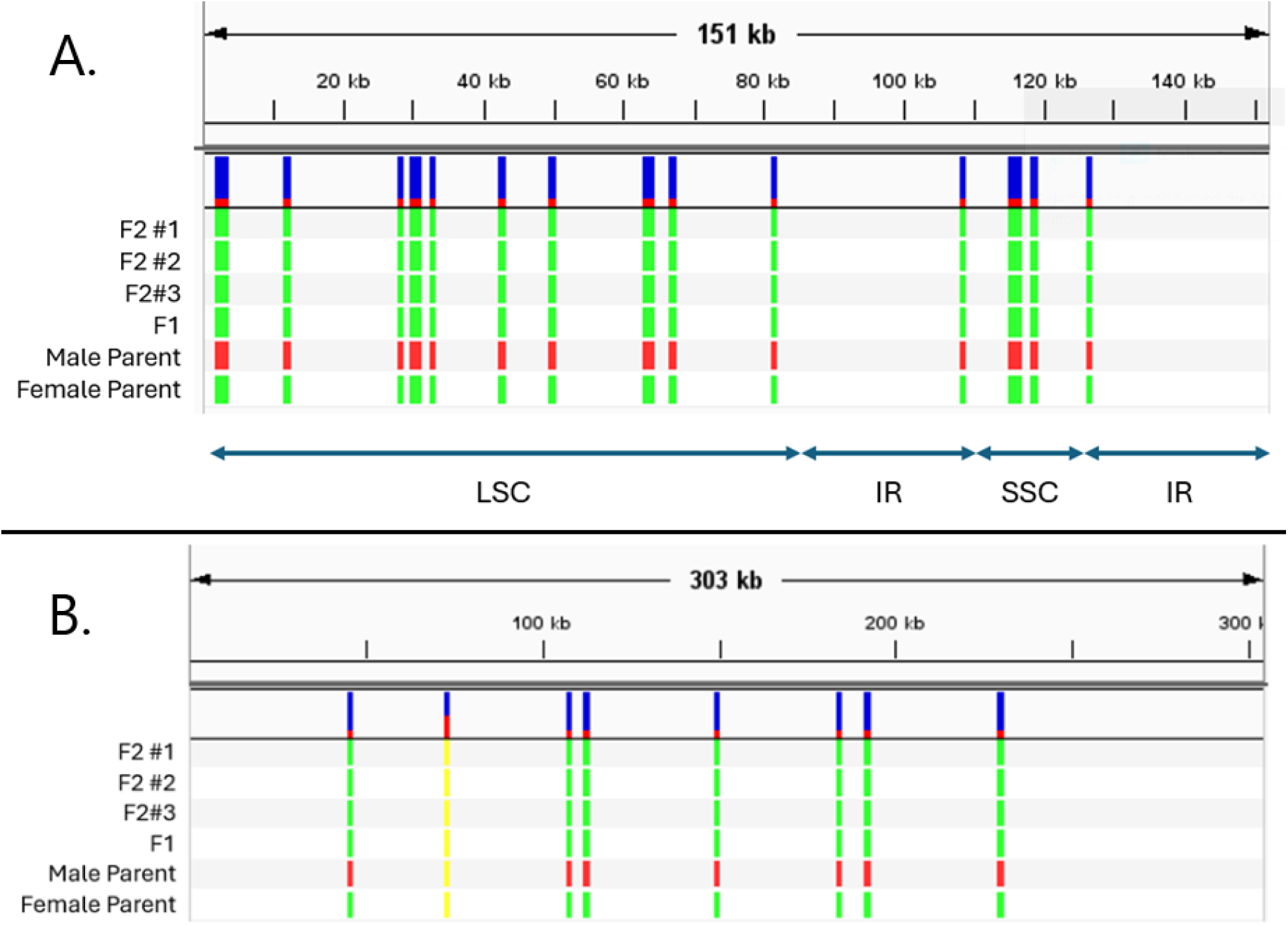
**A**. Chloroplastic ‘Path 2’ polymorphisms found among three F2 individuals, one F1 individual, and the maternal (’QC’) and paternal (’P’) parents in the cross that yielded the F1 individual. **B**. Mitochondrial polymorphisms found among three F2 individuals, one F1 individual, and the male and female within the cross that yielded the F1 individual, ‘QC’ and ‘P’ respectively. Interestingly one polymorphism shared across all individuals is a heterozygous A/C SNP.

To determine the transmission pattern of the mitochondrial genome in the ‘P’ x ‘QC’ cross, ‘QC’ short read alignment to the ‘P’ mitochondrial reference genome detected nine polymorphisms, of which eight were informative, with the uninformative polymorphism being a SNP for which all individuals had a non-reference nucleotide. Seven of those informative polymorphisms were SNPs, while the eighth was a 22 bp indel (Figure 5B, Supplemental Table 3). These data demonstrate that transmission of the mitochondrial genome in the ‘P’ x ‘QC’ cross was maternal.

### Chloroplast and Mitochondrial Genome Ancestries

A phylogeny based on *de novo* assembled chloroplast genomes from *B. vulgaris, C. quinoa, C. berlandieri* var. *macrocalycium, C. foggii,* and both the ‘P’ and ‘QC’ *C. ficifolium* accessions (Figure 6A) revealed a close affinity of the two *Chenopodium* allotetraploids with the A-genome diploid *C. foggii*, and a more distant relationship with B-genome diploid *C. ficifolium*. Similarly, mitochondrial reference-based assemblies utilizing the same set of germplasm (Figure 6 B) in a narrowly focused phylogeny detected the same pattern of affinities. These outcomes support the prior hypothesis (Maughan *et al*., 2019) that the organelle genomes of the AABB allotetraploids were both contributed by the AA ancestral diploid.

**Figure 6.**
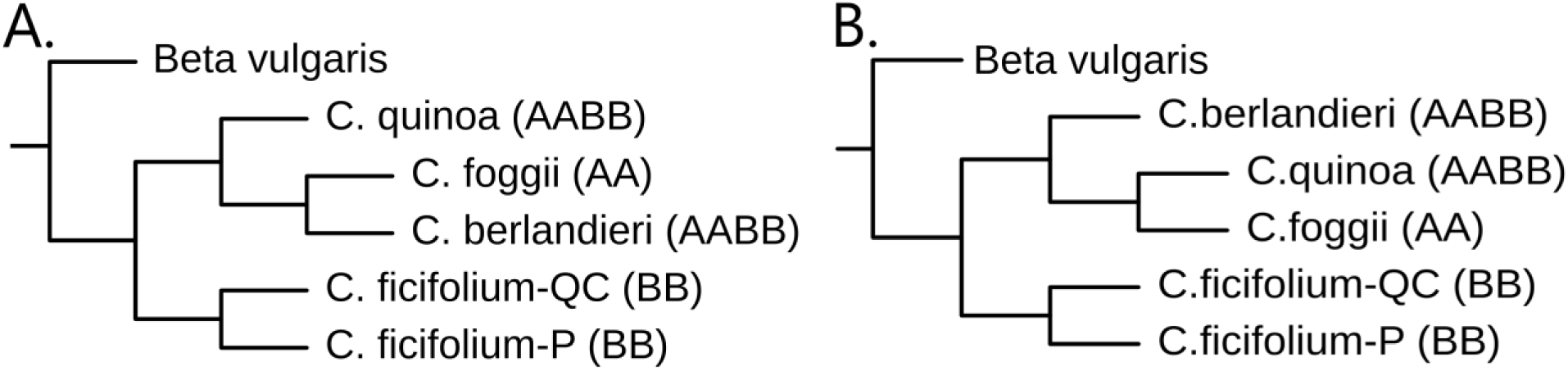
**A**. Phylogenetic tree illustrating chloroplast genome ancestral relationships among Chenopodium AA and BB diploids and AABB tetraploids in our study. Phylogenetic tree illustrating chloroplast genome ancestral relationships among Chenopodium AA and BB diploids and AABB tetraploids in our study. This phylogenetic tree is based on chloroplast assemblies from C. foggii, both C. ficifolium accessions, C. quinoa, C. berlandieri, and Beta vulgaris which is set as the outgroup. **B**. Phylogenetic tree illustrating mitochondrial genome ancestral relationships among Chenopodium AA and BB diploids and AABB tetraploids. This phylogenetic tree was constructed using reference guided assemblies with the C. quinoa mitogenome (NC_041093.1) selected as the reference genome. Beta vulgaris is set as the outgroup.

### Genome Wide Association Study

Of the five trait-specific GWAS analyses performed, two (branch angle and internode length) failed to detect genomic associations. The remaining three traits – days to flowering, plant height, and branch number – were found by GWAS to correlate with a single common region of chromosome Cf4 (Figure 7 A-C). Of these three traits, the correlated SNPs for branch number covered the largest region and fully contained those implicated in the plant height and days to flower analyses. The implicated region spans 7,020,561 bp between positions 70,765,247 and 77,785,808 on Cf4. Features within this region were pulled from the annotation file to create a list of gene names and GO terms associated with those genes. There are 770 annotated genes within the defined region. Of those, 132 were of unknown function. GO terms were assigned to 403 genes, including *FTL1*, which at position 74,807,401 – 74,811,571 is number 425 on the list of 770 genes, and is annotated as *Vernalization 3*. *FTL1* had one product classification identified by Interpro (Paysan-Lafosse *et al*., 2023), being a phosphatidylethanolamine-binding protein (IPR008914).

**Figure 7.**
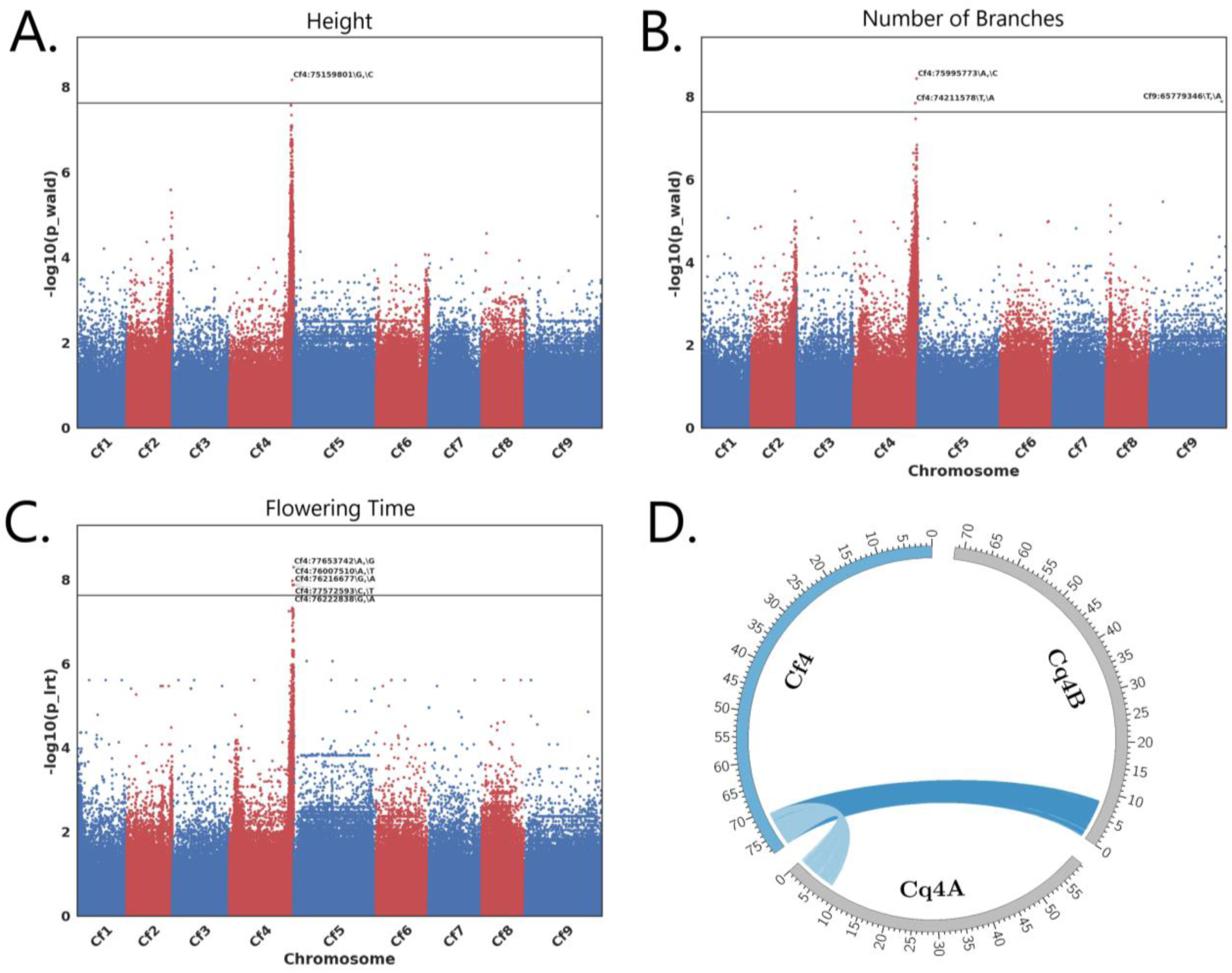
**A-C.** GWAS Manhattan plots indicating regions of the genome implicated in the control of plant height, number of branches, and flowering time. **D**. Synteny analysis of the ∼7 Mb region found in C. ficifolium Cf4 (Left) implicated by the genome wide association study as compared to C. quinoa chromosomes Cq4A and Cq4B as oriented in Rey et al., (2023). Both quinoa chromosomes Cq4A and Cq4B are in reverse orientation compared to C. ficifolium chromosome Cf4.

A microsynteny analysis focusing on the ∼7 Mb window implicated in this GWAS was performed to identify homeologs within both *C. quinoa* subgenomes and their locations (Figure 7 D). MCScanX identified nine collinear blocks of 10 or more genes from this region in *C. ficifolium* that also occur, but in reverse orientation, on quinoa chromosomes Cq4A and Cq4B of the Rey et al., (2023) assembly, as will be discussed later. The analysis of these data indicates that for the 770 genes found in *C. ficifolium* annotation file at this ∼7 Mb region, there were 581 homoeologous genes found within the *C. quinoa* A subgenome and 610 found within the B subgenome. In total, 107 of the *C. ficifolium* genes within this region are of unknown function and of those, only six have associated GO terms.

## Discussion

### Nuclear Genome

As described here the ChenoFicP_1.0 assembly provides a valuable component to the *C. ficifolium* diploid (BB) model system, as will be expanded upon in subsequent reports on ongoing investigations at the diploid and allotetraploid (AABB) levels in *Chenopodium*. Pseudochromosomes within this *C. ficifolium* assembly have been numbered to correspond with, and oriented to be syntenic and collinear with, the previously published *Beta vulgaris* (2n = 2x = 18) EL10.1 genome (McGrath *et al*., 2023), thereby following the example of Jarvis et al., (2017) in which the quinoa V1 assembly was similarly oriented. The fact that the quinoa V2 assembly (Rey *et al*., 2023) did not follow this orientation pattern opens the possibility for some confusion, especially with respect to chromosomes Cq1B, Cq4B, Cq5B, Cq7B and Cq8B, which are presented and numbered in reverse orientation in quinoa V2 as compared with quinoa V1 and beet EL10.1. We have followed the initial orientation approach in conjunction with the recently published *Chenopodium* pangenome analysis (Jaggi *et al*., 2024), which employs the original orientation pattern.

Beet was initially selected as the genome comparator for *Chenopodium* because it was the first species to have a published reference genome within the subfamily Chenopodioideae. While the diploid *Spinacia oleracea* (2n = 2x = 12) now has a published genome (Cai *et al*., 2021), its nuclear genome has only six chromosomes, making comparisons with it suboptimal for identifying syntenic relationships and orienting chromosomes of newly sequenced species. Genome assemblies have been reported for three other *Chenopodium* diploids: the AA diploids *C. pallidicaule* (Jarvis *et al*., 2017), *C. watsonii* (Young *et al*., 2023), and the BB diploid *C. suecicum* versions 1 (Jarvis et al., 2017) and 2 (Rey et al., 2023). However, to our knowledge, no gene-trait association studies have yet been conducted in these species.

The ChenoFicP_1.0 nine-pseudochromosome assembly size of ∼730 Mbp incorporates 96% of the 760 Mbp total assembly for the 781 contigs generated by Hifiasm. Cytometrically determined C-values of *C. ficifolium* ‘Portsmouth’ predict a genome size in the range of 821 to 831 Mbp (Neff, 2017). For comparison, the nine B-subgenome pseudochromosomes of the quinoa assembly sum to 670 Mbp in length (Rey *et al*., 2023), or approximately 8% smaller than the ChenoFicP_1.0 assembly. Similarly, the nine pseudochromosomes of the quinoa A subgenome sum to 531 Mbp (Rey *et al*., 2023), which is 3% smaller than the diploid *C. watsonii* 548 Mbp A-genome assembly (Young *et al*., 2023). An earlier, broad survey of *Chenopodium* germplasm reported C values of 893 Mbp for a Eurasian accession of *C. ficifolium*, 645 Mbp for *C. watsonii*, and 1.545 Gbp for *C. quinoa* (Mandák *et al*., 2016) Thus, all three of the aforementioned *Chenopodium* genome assemblies are somewhat smaller than the size predicted by cytometric assay. It has been speculated that this pattern of discrepancy may be due to incomplete assembly coverage of some repetitive regions (Young et al., 2023), or it could be due to a consistent technical artifact of flow cytometric analyses.

The ChenoFicP_1.0 genome contains 22,617 genes (total genes), of which 5,060 encode proteins that have yet to be characterized within the UniProtKB/Swiss-Prot database. Gene density is greatest in the distal regions of chromosomes (Figure 1 Ring A). For comparison, the total gene counts of the *C. quinoa* A and B subgenomes are 25,913 and 26,942 respectively (Rey *et al*., 2023), the AA diploid *C*. *watsonii* count was 30,325 (Young *et al*., 2023), and the BB diploid *C. suecicum* count was 29,702 (Rey *et al*., 2023). Compared with the ChenoFicP_1.0 assembly, the total number of annotated genes within the *C. quinoa* B subgenome is approximately 19.1 percent higher. Given that different gene count methodologies were used in these various studies, the observed differences in gene counts among subgenomes and species should be interpreted with caution.

RepeatModeler indicated that repetitive elements account for approximately 70 percent (510 Mbp) of the *C. ficifolium* genome. For comparison, the quinoa repetitive element content was approximately 65% for the total genome (Zou *et al*., 2017; Rey *et al*., 2023), and approximately 63% (421 Mbp) for the B subgenome, with the largest contributors being long terminal repeat retrotransposons representing 46 percent of the total genome size. Over half of the *C. ficifolium* repetitive elements are retrotransposons that cluster towards the center of chromosomes (Figure 1 Ring B). These data indicate that there has been some change in the repetitive element landscape in *C. ficifolium* and/or the *C. quinoa* B subgenome subsequent to the quinoa allotetraploid speciation event.

### Organelle Genome Assemblies

The generation of two different chloroplast genome assembly versions, designated Paths 1 and 2, was an unexpected outcome. Prior chloroplast genome assemblies in *Chenopodium*, including *C. quinoa* (Maughan *et al*., 2019; Gao *et al*., 2021) and *C. ficifolium* (Kim, Chung and Park, 2019), and in the broader Amaranthaceae family (Chaney *et al*., 2016; Dong *et al*., 2016; Kim, Chung and Park, 2019; Ding *et al*., 2021) including sugar beet (McGrath *et al*., 2023), have reported only a single chloroplast genome configuration, corresponding to our Path 2 assembly, although Kim et al. (2019b) mention the need to reposition the SSC and LSC regions of the quinoa accession ‘Real Blanca’ for their alignment procedure, indicating a disagreement in SSC and/or LSC orientation when compared to other assemblies. However, recent literature (Wang, Lanfear and Gaut, 2019) (and citations therein) has established that chloroplast genomes do indeed exist in two equally represented alternate configurations in many if not most plant species, as has now become evident in *Chenopodium*, and that the detection of these alternate forms is enhanced by employment of long read sequencing technologies such as PacBio HiFi and Nanopore. The very long and accurate reads now available from Nanopore technology will be of particular value in relation to mitochondrial genomes, the potential alternate forms of which are often challenging to assemble (Kozik *et al*., 2019). Given the known complexities of mitochondrial genome organization, the assembly configuration that we present here should be viewed as preliminary and subject to revision.

### Organelle Transmission Patterns

In the absence of evidence to the contrary, organelle genomes are generally assumed to be transmitted maternally in Angiosperms: however, exceptions have been reported (Davis *et al*., 2010), including paternal chloroplast transmission in some *Actinidia* hybrids (Testolin and Cipriani, 1997). Until very recently (Maughan *et al*., 2024), empirical documentation of modes of organelle inheritance in *Chenopodium* appears to have been limited to a single study involving atrazine resistance in *Chenopodium album* (Warwick and Black, 1980; Corriveau and Coleman, 1988; Harris and Ingram, 1991; Kolano *et al*., 2016). Our analysis agrees with this prior study in its finding of maternal transmission of both chloroplast and mitochondrial markers in our diploid level *C. ficifolium* cross. Similarly, Maughan et al. (2024) have documented the maternal transmission of the organelle genomes in several tetraploid level *Chenopodium* crosses.

### Organelle Genome Ancestries

Although *C. ficifolium* is a candidate B nuclear genome donor in the initial hybridization that formed quinoa 3.3 to 6.3 million years ago (Štorchová *et al*., 2015; Kolano *et al*., 2016; Jarvis *et al*., 2017; Maughan *et al*., 2019), prior analyses indicate that a BB diploid was not the ancestral donor of the allotetraploids’ organelle genomes (Maughan *et al*., 2019). Instead, that role has been assigned to the A nuclear genome donor (Maughan *et al*., 2019). That assignment is supported by our chloroplast and mitochondrial minimal phylogenies, which both place *C. ficifolium* as sister to a clade consisting of AABB tetraploids and an AA diploid.

It is considered likely that, in the initial AABB hybridization event, the ancestral AA diploid served as female and maternal organelle genome donor (Maughan *et al*., 2019). Documentation of such in our *C. ficifolium* cross and in tetraploid level crosses (Maughan *et al*., 2024) strengthens the thesis that the original hybridization event was of the form AA x BB, i.e., with the AA diploid ancestor serving as female. Final validation of this model will await the analysis of organelle genome transmission in reciprocal crosses between AA and BB diploids.

### Genome Wide Association Studies

The data produced from these genome-wide analyses indicate that, in the studied segregating population, variation in the agronomically important traits of days to flower, plant height, and branch number were predominantly, if not entirely, under the control of one or more genes within a single, 7 Mb region located towards the bottom of pseudochromosome Cf4 and containing the *FTL1* locus previously employed as a marker by Subedi et al. (2021). Our genome-wide analysis extends the findings of Subedi *et al*. (2021) by focusing attention on this Cf4 “hot spot” and the genes residing therein. Of note, the association of branch angle with a single SNP on Cf9 was not further investigated due to its presumed spurious nature. Manhattan plots will typically associate a number of SNPs within and surrounding a causative region, and this questionable SNP did not have any significantly associated markers adjacent to it, nor was it seen in the other two analyses investigating plant height or days to flower.

Gene ontology data submitted to Revigo indicated the presence of a myriad of genes responsible for controlling functions that are not intrinsically associated with these traits of interest; however, this region contains *Flowering Locus T-Like 1* (*FTL1*), which was automatically annotated as *Vernalization 3* (*VRN3*) within the GFF file. This gene is between positions 74,807,401 and 74,811,571, placing it centrally within the denoted significance region on Cf4. *FTL1* had one product classification identified by Interpro (Paysan-Lafosse *et al*., 2023), being a phosphatidylethanolamine-binding protein (IPR008914).

To further understand relationships between *C. ficifolium* and *C. quinoa,* and functionally utilize *C. ficifolium* as a model species, BlastP and MCScanX microsynteny analyses were performed to investigate relative gene locations across species and subgenomes (Figure 7D). When comparing the *C. ficifolium* region of interest to both of the quinoa subgenomes, all homologous genes in syntenic blocks of ten or more genes were found to reside in corresponding distal regions of quinoa chromosomes Cq4A and Cq4B, reflecting substantial gene order and location conservation within the respective genomic window. This analysis identified quinoa genes CQ035263 (A subgenome) and CQ019992 (B subgenome) within the *C. quinoa* V2 annotation (GFF3) file as being homologous to *C. ficifolium VRN3* (*FTL1*). The product of CQ035263 was queried with BlastP and has an 83 percent sequence identity match to XP_021772286.1, *C. quinoa FLOWERING LOCUS T-like*. Similarly, the product of gene CQ019992 is identical to XP_021734330.1. *C. quinoa FLOWERING LOCUS T-like* has a 97% protein identity with the annotated *C. ficifolium* gene Cfic_20093. The data produced using *C. ficifolium* as a model organism for *C. quinoa* via this pipeline has been demonstrated to be an efficacious way of establishing gene-trait relationships via GWAS and locating syntenic regions across species that may be worth investigating to understand agronomically relevant traits. Further investigation of genes within this region is required, perhaps in conjunction with more high-depth sequence data and additional plants to facilitate a fine-mapping study; however, these data are strongly indicative of one or more genes within this region as being implicated in the control of these traits. Further research, likely employing gene editing or transformation methods, is required to test this hypothesis and firmly establish this relationship.

The results of this study, in particular the synteny and collinearity analysis comparing *C. ficifolium* and quinoa ‘QQ74’ chromosomes, presents intriguing hints regarding the potential usefulness of figleaf goosefoot as a breeding resource for quinoa improvement. The primary structural variants differentiating the B genome chromosomes in these two species are three large pericentric inversions. However, the largest of these, on chromosome Cq3B, was a rearrangement present only in a subset of quinoa lines derived from lowland Chilean germplasm, with at least 80% of quinoa lines expected to have collinear Cq3B in comparison with figleaf goosefoot (Rey *et al*. 2023). The remaining two inversions, on Cq1B and Cq4B, appear to have breakpoints extending less than halfway out into their respective chromosome arms, such that they do not encompass recombinogenic regions (Maughan *et al*. 2024). Consequently, while triploid (AB_q_B_f_) figleaf goosefoot x quinoa hybrids are expected to be highly sterile, backcross hybrids might theoretically be recoverable and, if so, we might expect a high degree of pairing and recombination between their B genomes. This approach – crossing of the polyploid crop to its wild, diploid progenitor - has been successfully employed to introduce numerous resistance genes from the wild progenitor goatgrass (*Aegilops tauschii*, DD) into bread wheat (*Triticum aestivum*, AABBDD) while restricting recombination, and consequently undesirable linkage drag, to a single subgenome (Cox, Raupp and Gill, 1994).

## Conclusion

The development of high-quality annotated reference nuclear and chloroplast genomes and a preliminary mitochondrial genome for *C. ficifolium* serves as a foundational advance in the development of this species as a BB diploid model system for related allotetraploids of interest, with a primary focus on *C. quinoa* and *C. berlandieri.* Looking ahead, next steps in the development and implementation of this model system include: broadening the germplasm base via plant collection in diverse environments; establishing new hybrids and populations segregating for additional traits of interest including male sterility (Ward, 1998), and disease resistance, specifically downy mildew (Nolen *et al*., 2022); and the establishment of effective protocols for gene editing and *in vitro* culture and regeneration for both quinoa and its diploid model(s), with immediate focus on manipulation of *FTL1*. The fact that the organelle genomes present in allotetraploid quinoa were likely derived via maternal transmission from its AA diploid ancestor, not from its BB ancestor, points to the need for parallel development of an AA diploid model system in *Chenopodium*. Foundational steps in this direction have already been taken via the generation of genome assemblies for AA diploids *C. pallidicaule* (Mangelson *et al*., 2019) and *C. watsonii* (Young *et al*., 2023), and useful germplasm collection in *C. foggii* (Neff, Sullivan and Davis, 2018). As yet, to our knowledge, no genetic crosses or segregating progeny populations have yet been generated in an AA diploid, so this should be a top priority going forward. Finally, opportunity exists for breeding of quinoa with figleaf goosefoot as an avenue for gene introgression from the diploid to the tetraploid level.

## Abbreviations

AED: annotation edit distance
EST: expressed sequence tag
GO: gene ontology
GWAS: genome wide association study
HCGS: UNH Hubbard Center for Genome Studies
HMW: high molecular weight
MSA: multiple sequence alignment
ONT: Oxford Nanopore Technology
PC: pseudochromosome

## Acknowledgements

The research reported in this publication was partially funded by the New Hampshire Agricultural Experiment Station. This is Scientific Contribution Number 3032. This work is/was supported by awards to Thomas Davis, UNH, from the USDA National Institute of Food and Agriculture (Hatch) Project NH00678 (accession number 1019990), and by USDA NIFA project 110390. Ed Wilcox and the BYU DNASC (RRID: SCR_017781) provided PacBio sequencing services, as compensated by USDA NIFA project 110390.

## Conflicts of Interest

These authors have not declared any conflicts of interest.

## Data Availability

The reference genome and sequence datasets used to create it are available from NCBI using IDs SUB12919328 and SRR28959327, respectively. The reference genome can be found using the search term “UNH_ChenoFicP_1.0”. The reference assembly, annotation data in GFF3 format, and supplemental annotation files can be accessed via figshare.com: https://doi.org/10.6084/m9.figshare.c.7590620. Sequence data for F2 individuals used in GWAS analyses are hosted on NCBI: https://www.ncbi.nlm.nih.gov/bioproject/PRJNA1262536. Code used can be found on GitHub: https://github.com/UNH-DavisLab/Ficifolium-Genome-V1/

## Supplemental Tables

**Supplemental Table 1.** Individuals included in GWAS and phenotypic data used in analyses.

| Individual | Flowering Time | Height | No. Branches | Branch Angle | Internode Length |
| --- | --- | --- | --- | --- | --- |
| 4 | 23 | 46.5 | 13 | 71.9 | 3.6 |
| 5 | 25 | 38.1 | 17 | 68.2 | 2.2 |
| 6 | 32 | 64.8 | 27 | 61.6 | 2.4 |
| 7 | 25 | 43.4 | 14 | 69.4 | 3.1 |
| 8 | 21 | 36.6 | 15 | 58.9 | 2.4 |
| 9 | 23 | 46.7 | 13 | 63.3 | 3.6 |
| 10 | 27 | 52.1 | 14 | 68.8 | 3.7 |
| 11 | 35 | 72.4 | 26 | 78.2 | 2.8 |
| 12 | 23 | 55.6 | 19 | 75.4 | 2.9 |
| 13 | 21 | 45.7 | 15 | 79.5 | 3 |
| 14 | 39 | 74.9 | 27 | 75.5 | 2.8 |
| 15 | 23 | 35.1 | 12 | 63.3 | 2.9 |
| 16 | 21 | 36.3 | 14 | 72.8 | 2.6 |
| 17 | 23 | 50 | 16 | 66.6 | 3.1 |
| 18 | 21 | 30.5 | 15 | 64.4 | 2 |
| 9r2 | 18 | 54.7 | 19 | 71.5 | 2.9 |
| 10r2 | 20 | 50.7 | 17 | 71.9 | 3 |
| 11r2 | 22 | 60.6 | 23 | 73.5 | 2.6 |
| 12r2 | 22 | 40.5 | 18 | 76 | 2.3 |
| 13r2 | 22 | 59 | 19 | 72 | 3.1 |
| 14r2 | 29 | 72.8 | 29 | 63 | 2.5 |

**Supplemental Table 2.** Centromere positions and lengths as identified by quarTeT CentroMiner and telomere lengths and directions as identified by quarTeT TeloExplorer.

| ID | Centromere |  |  | Telomere |  |  |  |  |
| --- | --- | --- | --- | --- | --- | --- | --- | --- |
|  | Start | End | Length | Present | Upstream Length | Dir. | Downstream Length | Dir. |
| Cf1 | 58004969 | 58247898 | 242930 | left | 3668 | + | 0 |  |
| Cf2 | 79538569 | 79657117 | 118549 | right | 0 |  | 3941 | - |
| Cf3 | 49844598 | 51718346 | 1873749 | both | 9779 | + | 3549 | - |
| Cf4 | 39049495 | 40750452 | 1700958 | both | 20412 | + | 5019 | - |
| Cf5 | 41797451 | 42801417 | 1003967 | both | 17843 | + | 3731 | - |
| Cf6 | 6996291 | 7223985 | 227695 | both | 5397 | + | 4382 | - |
| Cf7 | 38832653 | 41378639 | 2545987 | both | 882 | - | 5054 | - |
| Cf8 | 34002546 | 34566367 | 563822 | both | 4480 | + | 5243 | - |
| Cf9 | 46653552 | 46799075 | 145524 | both | 3710 | + | 4753 | - |

**Supplemental Table 3.**
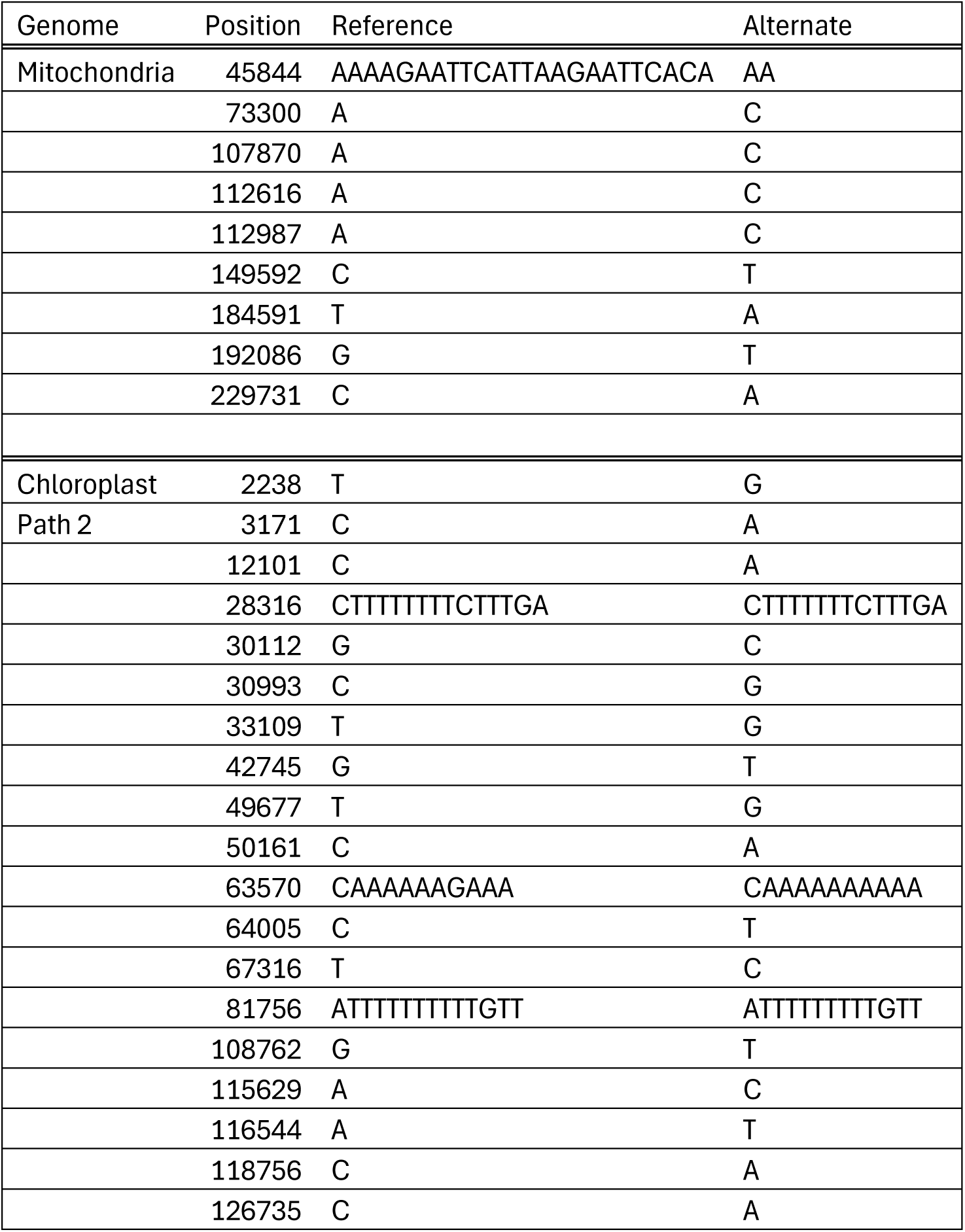
Positions and alleles of SNPs found in mitochondrial and chloroplast genomes used to establish maternal cytoplasmic inheritance.

## Notes

https://doi.org/10.6084/m9.figshare.c.7590620

https://www.ncbi.nlm.nih.gov/bioproject/PRJNA1101583

https://www.ncbi.nlm.nih.gov/sra/?term=SRR28959327

https://www.ncbi.nlm.nih.gov/sra/?term=SRR28959326

https://www.ncbi.nlm.nih.gov/bioproject/PRJNA1262536

https://github.com/UNH-DavisLab/Ficifolium-Genome-V1/

### Competing Interest Statement

The authors have declared no competing interest.

### Summary of Updates

Updated data availability section, revised methods regarding who initially assembled the genome used for analyses.

